# A Paradigm Shift in Walking, Sleep, and Exercise: Unique Effects on Blood Oxygen Saturation, Oxygen Diffusion, and Cellular Metabolism

**DOI:** 10.1101/2025.06.20.660810

**Authors:** P.A. “Tony” Gryffin, Q. Gu

## Abstract

Hypoxia underlies or complicates a wide range of chronic conditions, including cancer, arthritis, chronic pain, multiple sclerosis, stroke, chronic kidney disease, diabetes and more. Research is presented supporting indications that slower-paced exercises may develop states of relaxation, combined with enhanced respiration, which may trigger accelerated diffusion and facilitated oxygen use in the cells, stimulating cellular regeneration and healing. Metarobic theory is proposed as a physiological explanation for the benefits of slower-paced exercises, and as a good fit with aerobic and anaerobic categories of exercise. Metarobic theory posits that the momentary post activity drop in blood oxygen saturation (SpO2) following slow-paced exercises, and intermittently during sleep, may be the result of increased oxygen diffusion and metabolism. Metarobic effects may explain the non-aerobic health benefits of slower paces of walking, tai chi, qigong, and other slow-paced exercises, as well as the healing benefits of sleep. Research from the current study is presented, which supports that the momentary large drops in SpO2 ranging from 85% to 92% (*m=*89.2%±1.79) following slower-paced exercises, and periodically during sleep, may indicate a shift in the use and metabolism of oxygen. It is suggested that the momentary drop in SpO2 may follow an accelerated period of oxygen diffusion in response to hypoxic areas of the body, and a need for enhanced healing and cellular regeneration. In relation to sleep, this contrasts to current theory that lower levels of SpO2 during sleep result from more shallow respiration, due to the body needing less oxygen during sleep, resulting in hypoxemia. The end effect of slow-paced exercises and sleep may be to reverse or moderate hypoxia in the body through metarobic effects including enhanced oxygen diffusion, supporting healing and cellular regeneration.

## Introduction

The following article begins with a narrative of what led to the insights and research presented in this paper, followed by the methods, results, and discussion of the potential and limitations of the current research. My interest and understanding into the potential biological mechanisms which may underly many of the health benefits of slow-paced exercises, and the potential association with sleep, developed gradually over a period of years. Others have noted that slower-paced exercises, such as walking below 3.5 to 5 mph, do not raise the heart rate sufficiently to be considered aerobic forms of exercise for cardiovascular health..^1–5^

Chastin and colleagues noted in a systematic review of 72 studies that light intensity activities, such as slower paces of walking, are associated with beneficial acute and long-term lower risk of mortality. However, the authors stress that there is an absence of an established biological mechanism underlying benefits, which inhibits clinical recommendations.^1^ The authors note a need for research identifying the mechanisms underlying the benefits of light-intensity exercise.

Current modalities of exercise are divided into two primary categories: Aerobic exercise, which involves raising the heart rate to sufficient levels to generate cardiovascular benefits; and anaerobic exercise, which consist of activities which break down glucose for energy, such as weight lifting. As noted by Chastin and others above, slower paces of walking (non-speed walking) does not raise the heart rate to aerobic levels. And unless sped up to achieve an increased heart rate, neither do exercises such as forms of tai chi, qigong (breathing exercises), and yoga. This begs the question, what is happening in the body during slower-paced exercises, beyond benefits for stress reduction?^3,6–9^

My interest in this area started when students in my tai chi class at a community college reported various benefits for health at the end of each semester, including benefits for pain, feelings that tai chi and qigong enhanced their cancer treatment, and also benefits for arthritis and pain reported by my older students. I had suspicions that any potential benefits might be a result of enhanced oxygen use in the body, in part due to the literal definition of “qi.” In the tai chi community teachers attribute many of the benefits of these exercises to qi, generally translated rather vaguely as “vital energy.”^10,11^

However the literal definition of qi in most Chinese-English dictionaries is “air,” or oxygen.^11,12^It seemed possible that “qigong,” literally “breathing excellence” or “breathing exercise,” might enhance oxygen use in the body to stimulate healing. It is worth noting that although tai chi and qigong are generally listed as separate exercises, tai chi is actually one form of qigong. Qigong consists of a wide range of breathing exercises, and many of these are done independently, or as a “warm up” to tai chi.^13^

Based on the idea that enhanced oxygen use in the body might underly many of the benefits of these exercises, I began researching the role of oxygen and oxygen deficiency in various conditions. Hypoxia, or oxygen deficiency in the tissues, was a factor in every condition I searched, often with a large impact on the conditions development and/or outcome. This included cancer, cardiovascular disease, diabetes, kidney disease, chronic pain, arthritis, and immunity.^10^ Using a medical quality Nonin Go2 fingertip pulse oximeter, I began investigating potential changes in oxygen use during tai chi.^14,15^

Pulse oximeters, measuring oxygen levels in arterial blood, will typically not be impacted by cellar oxygen use, since venous blood is reoxygenated very efficiently by the lungs, usually back to normal levels above 94%.^16^ However, some conditions, such as chronic obstructive pulmonary disease (COPD), and extreme athletic performance,^17^ can result in low arterial blood oxygen saturation (SpO2) levels below 94%, as measured with a finger-tip pulse oximeter. This prompted the question, if exercises such as tai chi, being focused on oxygen in the body (qi), might also result in atypical changes in arterial blood oxygen saturation.

For my first investigation into this premise, I measured changes in SpO2 in 31 tai chi practitioners during practice.^14^ Minor increases in SpO2 were recorded during tai chi (*m=*1.29; p< 0.001). But what was particularly notable was a large drop which was observed following tai chi in one of the last participants. Just before the oximeter was removed, the SpO2 level dropped for just a few seconds to 88%, before returning to normal levels. Additional measurements during my own practice resulted in similar brief drops following tai chi and qigong practice, to as low as 82% in one instance.

This prompted a follow up study, conducting a series of 50 repeated measures in an experienced tai chi practitioner who also jogged regularly, to measure and document potential post-exercise drops in SpO2, during and for 5-minutes following tai chi, and also during and following running on a treadmill at 5 mph for comparison.^15^A similar slight increase was observed during tai chi (*m*=2.12±1.25%; p<0.001). But what was particularly notable was the large momentary post-tai chi drop in SpO2 levels (*m=*90.78±2%; p<0.001), ranging from 3 to over 10 seconds. No significant change was noted during or following running.

These results seemed potentially significant. Yet queries to various medical researchers yielded a general response that medical science already understood the effects of exercise on the body, and that the oximetry results were either in error, or were not of any significance. To better understand potential effects and to promote discussion and research in this area, and to address potential theory development, I conducted a review of over 200 studies on the role of hypoxia in various conditions, as well as studies on the potential benefits of tai chi and qigong for these conditions. I presented this research and theory development in a book which included an overview of what I have come to call metarobic theory, due to the potential effect of these exercises on oxygen metabolism, and as a good fit with aerobic and anaerobic categories of exercise.^10^

One particularly notable study found that by boosting oxygen diffusion around tumors in mice during radiation therapy, using an implantable micro-oxygen generator (IMOG), lower radiation doses were needed to achieve local tumor control, reducing radiation damage to the surrounding normal tissues, and potentially improving treatment outcome.^18^ In my own personal health, I suffered from what my doctors called celiac neuropathy. At that time I was practicing approximately 30 minutes of the traditional 108 yang stye tai chi form almost daily. I was curious if an additional focus on relaxation, by using small frame instead of medium frame tai chi, and using more relaxed arm postures in the non-dominant hand, as well as increasing dosage to 30 minutes 3-times a day, might enhance what I had come to call metarobic effects. The small frame and more relaxed format is similar to the modifications made by Cheng Man Ching, who was a noted teacher of tai chi for health.^19^

Within 3 months the neuropathy had completely cleared up. I reduced dosage to twice a day, or for a one hour session, which kept the neuropathy at bay. However, if I missed days I could feel the numbness start to return. So whatever was causing the neuropathy was still there, but the tai chi practice seemed to stimulate an enhanced state of healing and keep the neuropathy at bay. It is worth noting that even when I was just practicing tai chi fairly regularly once a day, though I had the numbness, I never experienced the pain associated with neuropathy, unless I missed several days of tai chi. It is also worth noting that benefits for pain is one of the most reported benefits of tai chi.^10,20^

Further reflection prompted the question, if any slow-paced exercise might enhance oxygen metabolism and healing, possibly by affecting hypoxia and oxygen diffusion. Much more research was needed, and my current position at the Mercer University School of Medicine provided further elements of support, including medical expertise and equipment. There was also the question of sleep. Stage 2 and stage 3 sleep, during which a large part of cellular regeneration and healing occurs, also involves a deep state of relaxation, and slower respiration. Could a similar physiological mechanism of action underly the healing benefits of sleep *and* slower paced exercises? Could the drop in arterial SpO2 be the result of a larger drop in venous blood oxygen levels (SvO2) below typical levels, due to enhanced oxygen use? As noted earlier, elite athletes experienced prolonged drops in SpO2 (91.9 ± 0.4%) due to a demand for oxygen in the muscles faster than oxygen can be replenished.^17^ This is discussed further, following the current findings presented below. The following results, from measurements of tai chi, qigong (breathing exercises), slower-paces of walking, and during sleep, compared to aerobic measurements, support that *something* different is going on in the body during these exercises. Possibly similar to changes in oxygen use during the healing stages of sleep, which may explain many of the health benefits of these exercises. Following is an overview of the methods, results, and discussion related to the current measurements.

## Methods

To obtain graphs and more accurate measurements of changes in blood oxygen saturation, a medical quality Nonin WristOx 3150 was used to record SpO2 levels. Unlike commercial oximeters which can miss brief changes in SpO2 level, the WristOx 3150 can record and prints out results in graph format for more accurate analyses. This device can record changes in SpO2 in 1 second intervals, with an accuracy of +/- 2%, including during challenging conditions which can include movement and low perfusion. This instrument is regularly used in military testing due to its durability and reduced error during movement. The motion tolerant software minimizes motion artifacts from being misinterpreted.^21,22^

An initial study was conducted at the Jewish Educational Alliance Fitness Center in Savannah GA during a summer scholars program through Mercer School of Medicine. Seven men and 3 women were included in the final fitness center group, ranging in age from 23 to 68 (*m=* 52.7 years). The participants included in the fitness center were in good health. One participant had recently completed an iron man competition (participant 2, Male, age 23). One participant (male, age 77) had been diagnosed with prostate cancer three months previously and was excluded from the fitness center group, in order to obtain a more accurate comparison of healthy individuals. Participants at the fitness center walked on a treadmill at 1.5 mph for 20 minutes, followed by 5 minutes of rest. Data was continuously recorded by the oximeter for download and evaluation.

As an exploratory study, additional measurements were taken of members of a tai chi group for comparison (see table 1). All members of this group noted a chronic condition, except for one member who did not share any chronic conditions on the background form. Participants in the tai chi group were measured while walking (at 1.5 mph), and during the practice of Yang style tai chi and the baduanjin qigong exercise, as well as for 5 minutes after each exercise while at rest. Measurements were conducted over a series of Saturday mornings during the groups regular meeting time of 9 am to 10 am. Additional measurements were also taken during sleep and moderate intensity aerobic exercise (using an exercise bike) in the lead researcher for comparison. Since heart rate would not return to resting levels within 5 minutes following using an exercise bike at moderate aerobic intensity, measurements were taken for 20 minute following the exercise period.

**Table 1.** Participants in the tai chi group.

| Participants in Tai Chi Group |  |  |  |
| --- | --- | --- | --- |
| Participant | Age | Sex | Condition |
| 1 | 66 | M | Type 2 Diabetes, Sleep Apnea |
| 2 | 55 | M | Recovering from Hernia Surgery |
| 3 | 68 | M | High Blood Pressure, Asthma |
| 4 | 71 | F | Chronic Kidney Disease, High Blood Pressure, High Cholesterol |
| 5 | 65 | M | High Blood Pressure |
| 6 | 69 | F | None Reported |
| 7 | 54 | F | Chronic Cytomegalovirus (CMV) Disease |
| 8 | 63 | F | Abdominal Cysts |
| 9 | 50 | F | High Blood Pressure, Lupus |
| 10 | 62 | M | Peripheral Neuropathy |

## Results

The graphs of the healthy participants in the fitness center group (*n*=10) did not show a clearly distinct drop in SpO2. However the older participant with prostate cancer (excluded from the fitness center group) did show a momentary marked drop in post-SpO2 down to 86%, before returning to normal levels. See figure 1. There were no clear drops in the other participants, except a small drop which was still within normal levels in participant 2 (See figure 2).

**Figure 1.**
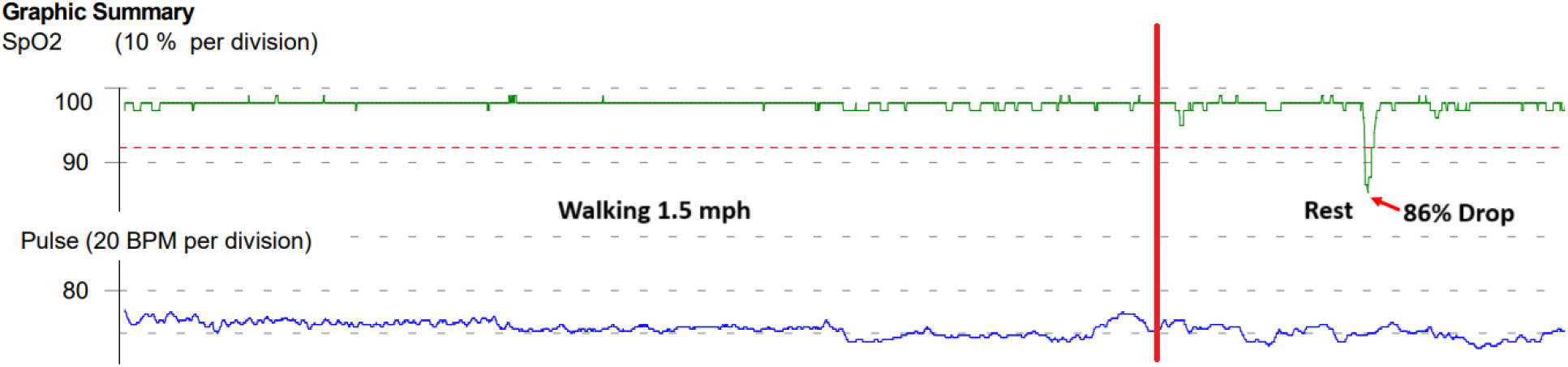
Line graph of SpO2 and heart rate of the older participant with prostate cancer, which showed a large distinct drop to 86% before returning to normal levels.

**Figure 2.**
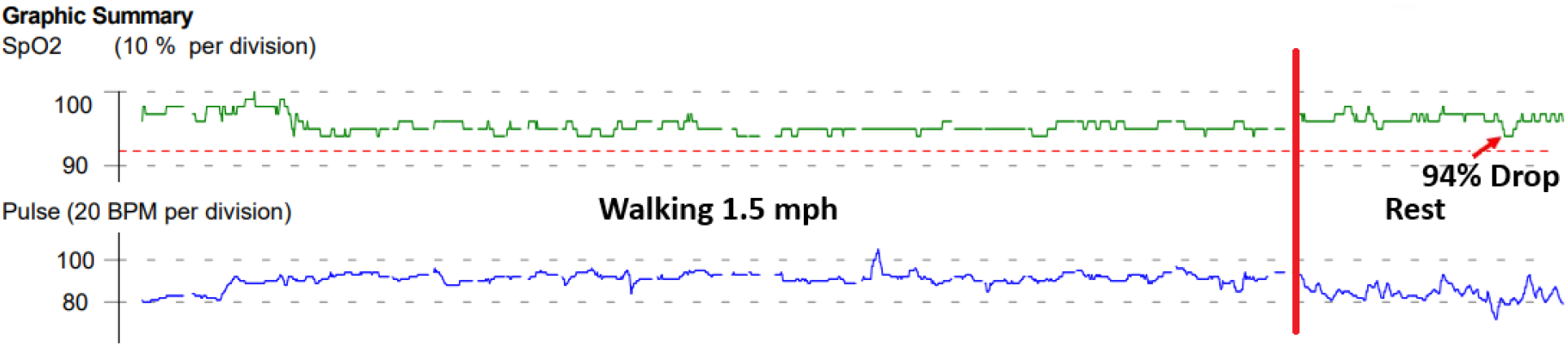
Participant 2 in the fitness center group experienced a potential distinct drop in SpO2 to 94%. See the discussion section for implications.

The measurements of the tai chi group yielded similar results to the older participant with prostate cancer, during walking, tai chi, and baduanjin qigong. Measurements taken of the primary author during sleep also exhibited period drops in SpO2 (see figure 7). However, during moderate aerobic activity no distinct post-activity drop in SpO2 occurred (see figure 6). Refer to table 1 for results of the fitness center group, the tai chi group, and the primary author. See figures 3 to 5 for sample graphs of slow-paced walking, tai chi, and qigong, and figure 6 for moderate aerobic activity (exercise bike), and figure 7 for sample sleep measurements.

**Figure 3.**
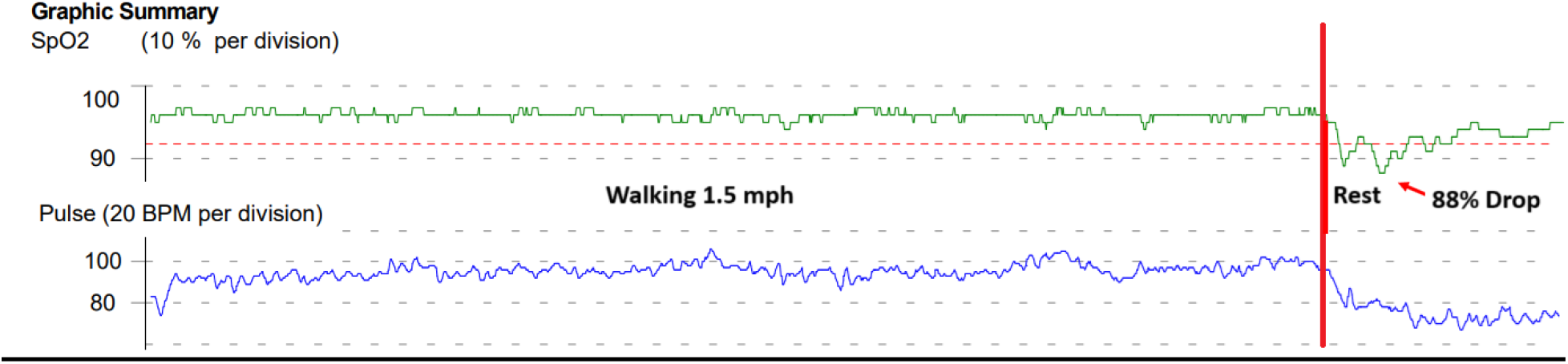
Measurements of participant 4 from the tai chi group during and following walking at 1.5 MPH.

**Figure 4.**
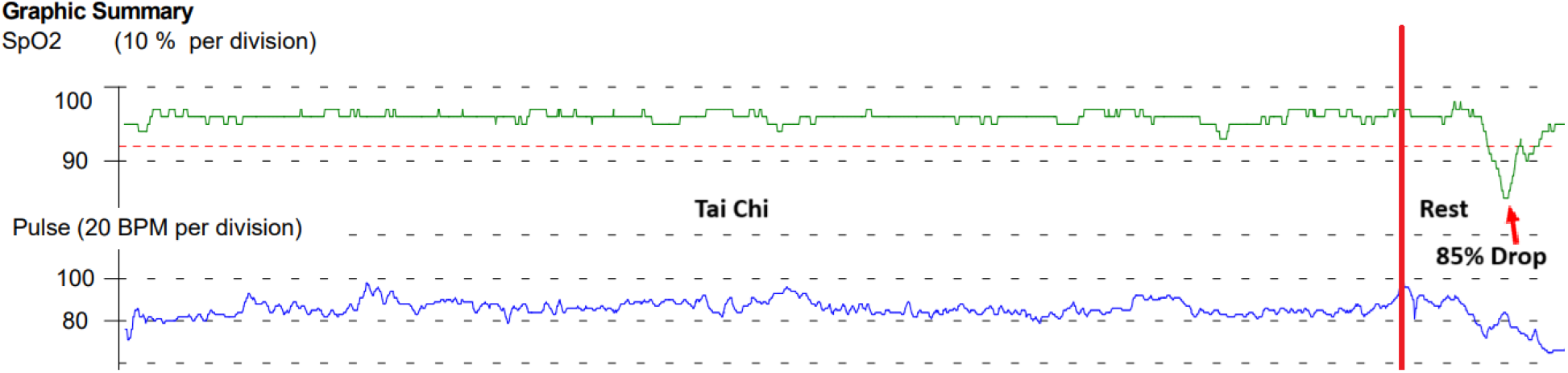
Measurement of participant 2 from the tai chi group during and following tai chi.

**Figure 5.**
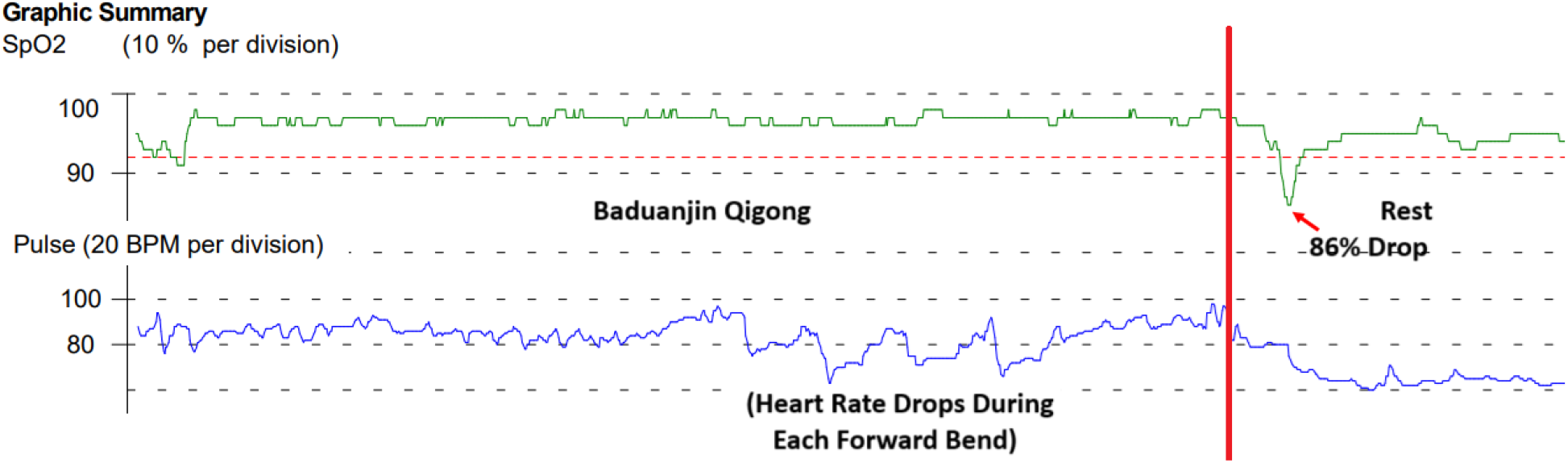
Measurement of participant 9 from the tai chi group during and following qigong (baduanjin).

**Figure 6.**
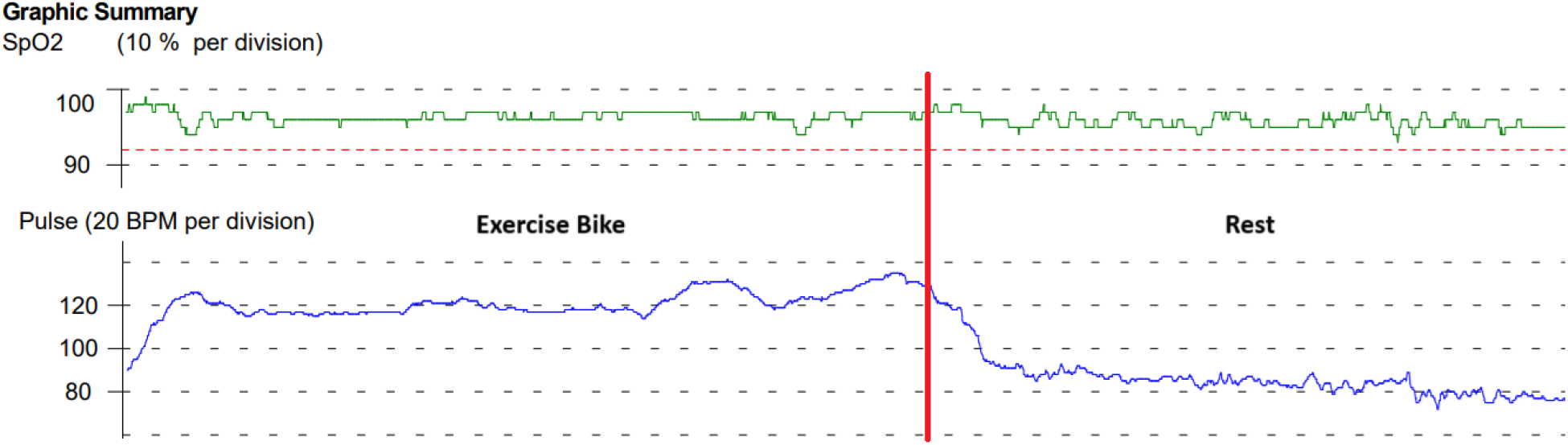
Measurement of the lead researcher during and following moderate intensity exercise on an exercise bike.

**Figure 7.**
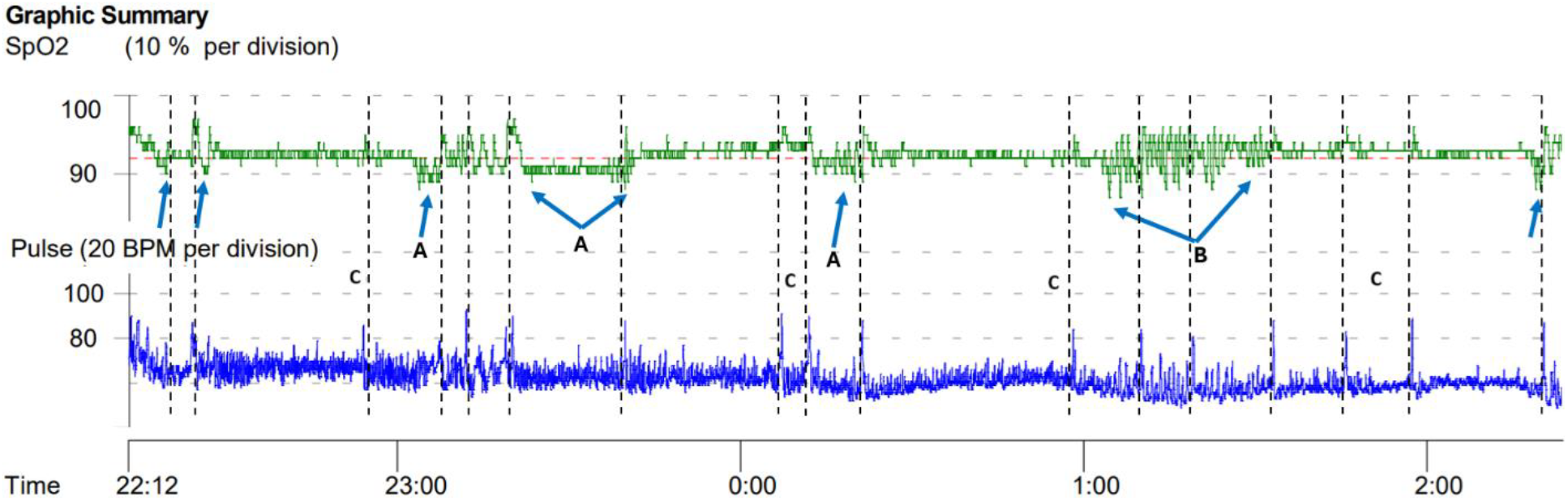
Measurement of a 4 hour sleep period for the lead researcher. Measurements which showed periodic drops in SpO2 to below 90% seemed to corresponded to some degree to an increase in heart rate to above 80 bpm, with a few exceptions (C), which may indicate a potential shift in metabolism and/or stage of sleep. Segments marked (A) included prolonged periods of lower SpO2, which may be due to a particular demand for oxygen. Segment (B) demonstrated a particularly rapid swing between lows and highs in SpO2 for approximately 25 minutes. See the discussion section for implications. In this series, Spo2 dropped to as low as 87%, with a mean SpO2 of 93.2%. Other measurements showed a similar pattern. See table 2 for mean scores during sleep.

## Discussion

Hypoxia underlies or complicates a wide range of chronic conditions, including arthritis, cancer, chronic kidney disease, diabetes, multiple sclerosis, stroke and more.^10,23^ Hypoxia consists of lower levels of oxygen content compared to normal states in organs, specific tissues, or cell types.^23^ Hypoxemia, low levels of oxygen in the blood, can be affected by blood flow to the lungs (perfusion), ventilation or airflow to the alveoli, and gas exchange through diffusion.^24,25^ The latter is particularly relevant to the current study and theory development, since enhanced diffusion may affect SpO2 levels. As noted early, pulse oximeters measure arterial as opposed to venous oxygen content. However, if venous blood oxygen saturation falls below a certain point, the lungs may not restore oxygen levels to normal levels.

**Table 2:** Results from the follow up measurements of the tai chi group* (Tai Chi, Qigong, Waking), compared to the healthy participants in the fitness center group** (Healthy Walking), and to sleep and moderate biking in the lead researcher.***

| Change in SpO<sub>2</sub> |  |  |  |
| --- | --- | --- | --- |
| Activity | SpO<sub>2</sub> During Activity | SpO<sub>2</sub> After Activity | *p*-Value |
| **Tai Chi\*** | 96.25 ± 0.98 | 89.30 ± 1.70 | <0.001 |
| **Qigong\*** | 96.60 ± 0.74 | 88.40 ± 1.43 | <0.001 |
| **Slow Walking\*** | 97.20 ± 0.95 | 89.90 ± 2.02 | <0.001 |
| **Healthy Walking\*\*** | 96.60 ± 1.37 | 96.80 ± 1.21 | 0.269 |
| **Sleep\*\*\*** | 93.90 ± 0.46 | 88.20 ± 1.62 | <0.001 |
| **Mod. Bike\*\*\*** | 95.80 ± 0.89 | 95.95 ± 0.76 | 0.468 |

The metarobic theory proposed posits that states of relaxation and slower respiration which can be engendered by slower-paced exercises may enhance oxygen diffusion and use in the cells, triggering a metabolic state which enhances healing and cellular regeneration. A metarobic effect may also occur during other activities as well, such as during sleep and sitting meditation (discussed below). See figure 8 for a visual representation of a potential metarobic effect.

**Figure 8.**
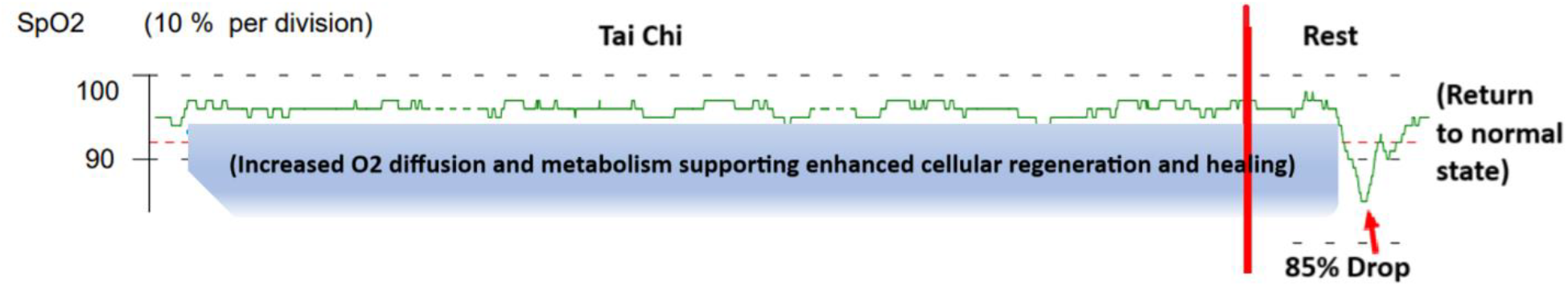
Metarobic theory posits that the momentary post-drop in oxygen levels may be the result a shift back to normal metabolism. Enhanced diffusion and oxygen metabolism may support cellular regeneration and healing. Metarobic effects may explain the non-aerobic benefits of slower paces of walking, tai chi, qigong, and other slow-paced exercises, as well as the healing benefits of sleep. The above shows the momentary drop in SpO2 to 85%, following 20 minutes of tai chi from figure 4. Similar drops occurred during walking, qigong, and sleep. See figures 1, 3, 5 and 7.

During the practice of tai chi and qigong, which are consciously focused on relaxation and the breath, it is common to feel a tingling in the fingers, hands, and sometimes extremities. Similar to when a limb falls asleep due to a lack of circulation, but pleasant instead of painful. It is suggested that this pleasant tingling is a result of enhanced oxygen diffusion through the cells.

Generally, when SpO2 levels drop below a certain threshold (typically considered to be around 92%), it indicates that the body is not adequately oxygenated, signifying hypoxia/hypoxemia.^26^ However, in the case of the slower-paced exercises measured in this study, and in previous research,^15^ the timing and short duration of these drops, sometimes to as low as 84% for a brief moment before returning to normal levels, suggests a state other than hypoxia or hypoxemia.

Oxygen and related signal transduction pathways play an important role in controlling cell proliferation during the development of many tissues, including the nervous system. Hurst and colleagues noted that oxygen levels in the cells, in relation to hypoxia, can affect cellular metabolism and physiological function, through various mechanisms that underlie oxygen sensing.^27^ It was observed that cells can sense and adapt to changes in oxygen availability, which can also regulate the activity of genes in response to variation to oxygen level. It is suggested that the relaxed state coupled with slower respiration may affect oxygen sensing mechanisms related to the increase in oxygen diffusion, stimulating enhanced cell metabolism, and supporting healing and cellular function. Other similarities between sleep and relaxation exercises such as tai chi, is a reduction in blood pressure.(Elkhenany et al.,2018; Yeh et al., 2008)

That the individuals in the fitness center group with no reported health conditions did not experience a clear drop in SpO2 following walking raises interesting questions. Being within the normal range of SpO2 any drops may not be particularly noticeable. Participant 2 in the fitness center group did have a potentially clear drop in SpO2 to 94% (figure 2), but not to the degree of the other participants in the tai chi group, nor the participant with prostate cancer. Any drops in SpO2 which are not below 92% may not be as noticeable. If the level of the momentary post activity drop in SpO2 is tied to the level of health of the person, healthier individuals may experience less of a drop in SpO2 compared to individuals with various chronic conditions. That the participants who experienced the large post-activity momentary drop in SpO2 were all older adults begs the question if aging in general places more demands on a need for ongoing cellular regeneration. Or is age independent of the chronic conditions which the participants in this group reported? And does level of relaxation play a role? The participants in the tai chi group may have experienced larger drops following slower-paces of walking due to having learned to function in a more relaxed state from their tai chi practice, whereas the participants in the fitness center group may maintain a level of tension even when walking at 1.5 mph. Indeed, one of the fitness center participants noted how difficult it was to force themselves to keep to a slow pace.

The primary premise being considered as metarobic theory, is that the momentary drops in SpO2 following slower-paced exercises, and periodically during sleep, may be a result in a shift in the use and metabolism of oxygen resulting from a relaxed state, potentially in response to hypoxic areas of the body, and a need for enhanced healing and cellular regeneration. In relation to sleep, this contrasts to current theory that lower levels of SpO2 during sleep result from more shallow respiration, due to the body needing less oxygen during sleep, resulting in hypoxemia. Based on figure 7 above, periodic drops in SpO2 below 92% may be consistent with the transition between stages of sleep, and may be a result of a similar physiological effect underlying the drops in SpO2 following slow-paced exercises, particularly those which have an element of relaxation.

In a review by Elkehenany and others, it was noted that the healing process is enhanced during sleep. However, the mechanism which underlies faster healing during sleep has not been studied to any satisfactory extent.^28^ The authors note that the literature states benefits of sleep may be due to mild hypoxic states, which Elkehenany notes does not make sense.^28^

The literature considers mild hypoxia during normal sleep, as a factor which may enhance the survival, proliferation, and function of stem cells. But they do not state the timing of the occurrence of the states of mild hypoxia.^28,29^ One current explanation for why hypoxia is needed for the maintenance of a stem cell pool is that mild to low levels of oxygen might minimize damage caused by oxidation.^29,30^ On the other hand, this theory is not universally accepted, since hypoxia has also been reported as increasing rather than reducing oxidative stress.^29,31^So an alternative explanation or hypotheses may be metarobic theory. Currently most literature in the area of sleep states that these brief periodic drops are random and meaningless, and that low levels of SpO2 in those without breathing related illnesses is due to a shift to shallow breathing, and lower availability of oxygen in the lungs.^28^

On the other hand, ongoing levels of mild hypoxia, at or around 92%, may be the result of increased oxygen demand for cellular regeneration and healing, rather than as a result of reduced oxygen intake during respiration. Metarobic theory suggests that mild hypoxia during sleep may be the result of healing and cellular regeneration, rather than the cause of healing and cellular regeneration.

In a discussion with Dr. Kelsey Joyce with the University of Birmingham, regarding his studies of nocturnal pulse oximetry and acute mountain sickness, Dr. Joyce was able to confirm that he saw similar periodic drops in SpO2 during sleep to those in figure 7. Periodic drops in SpO2 can also be seen intermittently during the prolonged drops in those with COPD, which according to metarobic theory, may indicate the body’s ongoing attempt at healing, even with the reduced oxygen content in the blood (figure 9).

**Figure 9.**
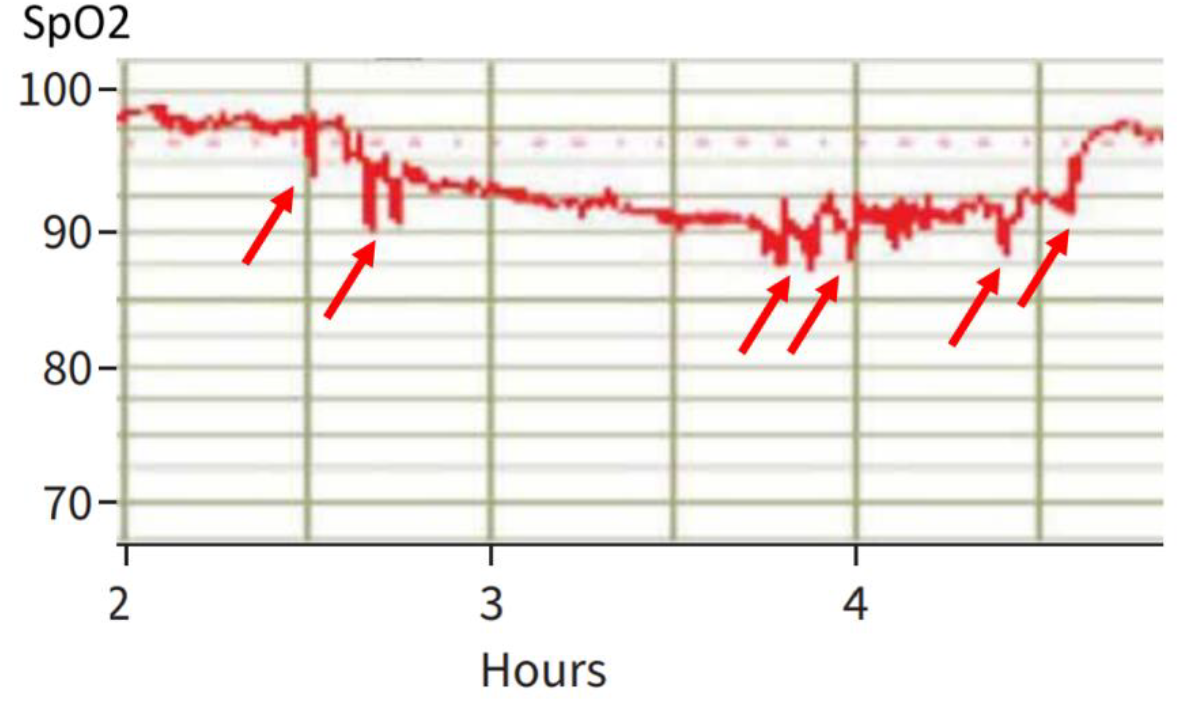
Overnight oximetry pattern segment consistent with COPD. The arrows were added by the author to indicate potential drops in SpO2 which may parallel similar drops during sleep in noted in figure 7, and following periods of slower-paced exercises.^32^

In the current study, breathing rate was measured with a Go Direct Respiration Belt. During tai chi, respiration slowed down from the 12 to 20 breaths per minute typical in healthy adults during rest,^33^to as low as 6 to 10 deep breaths per minute during tai chi, which is supported in the literature.^34,35^ Breathing rate was also observed while walking at 1.5 mph in the current study, to fall between 10 to 12 breaths per minute. Stages 2 and 3 of sleep result in a similar reduction of breathing rate.^36^

The difference between sleep and slower-paced exercises, related to depth of respiration (involving full abdominal and chest expansion and contraction), may explain why SpO2 increased slightly during the actual practice of slower paced exercises, but falls slightly during sleep, before experiencing the larger momentary drop in SpO2. The generally lower levels of SpO2 during sleep seen in figure 7 may be the result of less overall demand for oxygen by the body during sleep, possibly due to a reduced ongoing need by the large muscle groups which occur during wakefulness. During sleep there is a reduction in skeletal muscle blood flow.^28,37^ Under awake resting conditions, oxygen extraction in the muscle tissues ranges between 20% and 40%, while during heavy exercise this demand can increase to 70–80% of the oxygen carried by the blood.^38^ The relaxed state of slow-paced exercise and sleep may reduce the muscular need for oxygen, and enhance oxygen use by the rest of the cells of the body. Particularly in the organs, which are more prone to disease. Cancer development is very rare in skeletal muscle tissue.^39^ One reason relates to the larger demand for oxygen, and increased blood flow and perfusion in skeletal muscle tissue. Sleep and relaxation focused exercises may lead to an increase in O2 delivery to the organs, compared to the musculature during more active exercises and during waking and active hours.

A particularly interesting study found that during lower intensity exercises such as walking, patients with COPD actually had greater declines in oxygen saturation than when performing a higher intensity exercise such as cycling.^40^ The researchers noted that desaturation of approximately 7% occurs during an incremental treadmill test, compared to 3% during an incremental cycling test. Two theories have been proposed. One is that the increased drive to breathe is due to an earlier onset of metabolic acidosis during cycling, which led to higher ventilation. However, the researchers also proposed that the differences might be due to greater ventilation-perfusion mismatching with walking than with cycling. The researchers note that these two hypotheses are conflicting. As an alternative hypotheses, Mahler and colleagues note that their data supports that for those with COPD, enhanced alveolar ventilation and consequent higher partial pressure of oxygen (PaO2) may minimize the decrease in PaO2 and oxygen saturation in arterial blood during cycling compared to walking. The authors note that the initial increase in PaO2 during cycling occurred prior to the anerobic threshold, suggesting a neurogenic process possibly originating in the receptors of exercising muscles. The authors also suggested that the increased physiological demands of the leg muscles during cycling was a subsequent ventilatory stimulus.

However, as noted above, large muscles create a greater demand for oxygen in the body. Yet in the COPD group in the above study, walking, a less muscular demanding exercise, actually resulted in lower levels of SpO2 than during cycling. Could the higher level of desaturation be occurring during walking be due to greater oxygen diffusion and a metarobic effect, resulting from less muscle demands? See figure 9 for indications of periodic momentary drops during sleep in a patient with COPD, which may support a metarobic premise. The more relaxed state of the muscles might allow greater oxygen diffusion and cellular respiration, prompting greater O2 use for healing and cellular regeneration needed by those with COPD.

It is also worth noting that sitting meditation focuses on a relaxed stable posture and also focuses on relaxed breathing. Sitting meditation has been associated with a range of physical health benefits, including reducing oxidative stress, enhance telomere length, and associated cellular aging, however, how meditation affects these factors is unclear.^41^ Metarobic theory may offer one explanation.

Various limitations and cautions are worth noting. The first being the small number of participants, which may limit the ability to generalize. That the participants in the tai chi group were all older adults, with all but one identifying a chronic condition, is also a limitation in this study. Another limitation is that all of the older adults, except for the participant with prostate cancer excluded from the fitness group, practiced tai chi. It is possible that practice of tai chi enhances an ability to relax, which may result in contrasting effects with the fitness center group who were not tai chi practitioners. It is recommended to replicate these measurements with a larger number of participants in various age cohorts, in those who do not do tai chi, as well as for a wider range of practices, such as sitting meditation.

Also needing to be considered is the potential of undocumented COPD or other respiratory conditions, which may affect results. Another limitation is the lack of sleep graphs documenting SpO2 in the literature for healthy adults. In speaking with different sleep labs, periodic drops in SpO2 in patients without COPD or other breathing exercises occurred, but they were not able to share the graphs.

The observations presented in this research presents potential relationships, but these relationships may be independent of actual cause and effect, and without meaning. As an exploratory study, possible associations have been presented which might explain the post slow-paced exercise drop in SpO2 in relation to the benefits reported for these exercises. This is necessary at this stage, but may also result in bias and inclusion of potential associations where none actually exist. Future research will confirm, deny, or provide alternate associations, towards enhancing the effectiveness and practice, research and promotion of these exercises, as well as a better understanding of any implications in relation to the shifts and periodic drops in SpO2 during sleep.

One last and potentially particularly important point. Since Sp02 dropped significantly in the current study in older adults with a range of chronic conditions, could SpO2 measurements during sleep and slower paces of exercise be used to detect potential chronic conditions or illness? It might be worth conducting a range of medical evaluations in those without a chronic condition, yet who exhibit a large post slow activity drop in SpO2, to see if these drops correspond to an unknown illness or condition. Carlson and others note that declines during sleep in regional cerebral oxygen saturation (rcSO2) in the old from resting baseline may help identify individuals who are at risk for cognitive decline.^42^

It is also important to note that for overall health, aerobic and resistance exercises are still key for cardiovascular health, and are perhaps the most important forms of exercise for health and longevity.^4,43^ However, for older adults and those with chronic conditions who have an inability to engage in vigorous exercise, slower-paced exercises with distinct and measurable effects and benefits for chronic conditions may offer a valuable supplement for health. Having an understanding of the physiological mechanisms underlying benefits for health may help promote tai chi, qigong, and other slow-paced exercises such as walking, similar to the effect that an understanding of the aerobic effects of fast-paced exercises had on popularizing running and other forms of aerobic exercise.^44,45^

## Notes

### Competing Interest Statement

The authors have declared no competing interest.

## REFERENCES

1. Chastin SFM, De Craemer M, De Cocker K, et al. How does light-intensity physical activity associate with adult cardiometabolic health and mortality? Systematic review with meta-analysis of experimental and observational studies. Br J Sports Med. 2019;53(6):370–376. doi:10.1136/bjsports-2017-097563

2. American College of Sports Medicine (ACSM). Starting a Walking Program.; 2011. Accessed August 26, 2024. https://www.acsm.org/docs/default-source/files-for-resource-library/starting-a-walking-program.pdf

3. Zuhl M. Tips for Monitoring Aerobic Exercise Intensity.; 2020. Accessed August 26, 2024. https://www.acsm.org/docs/default-source/files-for-resource-library/exercise-intensity-infographic.pdf

4. Liguori G. American College of Sports Medicine. (2000). ACSM’s Guidelines for Exercise Testing and Prescription. 11th ed. Lippincott Williams & Wilkins,; 2021.

5. Myers J. Exercise and Cardiovascular Health. Circulation. 2003;107(1). doi:10.1161/01.CIR.0000048890.59383.8D

6. Kelly P, Williamson C, Niven AG, Hunter R, Mutrie N, Richards J. Walking on sunshine: scoping review of the evidence for walking and mental health. Br J Sports Med. 2018;52(12):800–806. doi:10.1136/bjsports-2017-098827

7. Yeh GY, Wang C, Wayne PM, Phillips R. Tai Chi Exercise for Patients With Cardiovascular Conditions and Risk Factors. J Cardiopulm Rehabil Prev. 2009;29(3):152–160. doi:10.1097/HCR.0b013e3181a33379

8. Yeh GY, Wang C, Wayne PM, Phillips RS. The Effect of Tai Chi Exercise on Blood Pressure: A Systematic Review. Prev Cardiol. 2008;11(2):82–89. doi:10.1111/j.1751-7141.2008.07565.x

9. Li X, Chang P, Wu M, et al. Effect of Tai Chi vs Aerobic Exercise on Blood Pressure in Patients With Prehypertension. JAMA Netw Open. 2024;7(2):e2354937. doi:10.1001/jamanetworkopen.2023.54937

10. Gryffin P. Mindful Exercise: Metarobics, Healing, and the Power of Tai Chi: YMAA Publications; 2018.

11. Zhang YH RK. A Brief History of Qi. Paradigm Publications; 2001.

12. The Oxford Chinese Dictionary: English-Chinese, Chinese-English.. Oxford University Press.; 2010.

13. National Institutes of Health NC for C and IH. Qigong: What You Need To Know. https://www.nccih.nih.gov/health/qigong-what-you-need-to-know.

14. Gryffin P. Qi: Implications for a New Paradigm of Exercise. Integrative Medicine. 2013;12(1):36–40.

15. Gryffin PA, Diaz RE. Effects of Tai Chi and running on blood oxygen saturation: a pilot study. J Complement Integr Med. 2021;18(4):821–825. doi:10.1515/jcim-2020-0306

16. Torp KD MPPE et al. Pulse Oximetry. https://www.ncbi.nlm.nih.gov/books/NBK470348/.

17. Galy O, Hue O, Chamari K, Boussana A, Chaouachi A, Préfaut C. Influence of Performance Level on Exercise-Induced Arterial Hypoxemia During Prolonged and Successive Exercise in Triathletes. Int J Sports Physiol Perform. 2008;3(4):482–500. doi:10.1123/ijspp.3.4.482

18. Cao N, Song SH, Maleki T, et al. Radiosensitizing Pancreatic Cancer Xenografts by an Implantable Micro-Oxygen Generator. Radiat Res. 2016;185(4):431. doi:10.1667/RR14149.1

19. Gibbs T. Cheng Tzu: Master of the Five Excellences A Life Biography of Cheng Man Ching by. https://web.archive.org/web/20150224133506/ http://sinobarr.com/cheng/cheng_life_bio.htm.

20. Kong LJ, Lauche R, Klose P, et al. Tai Chi for Chronic Pain Conditions: A Systematic Review and Meta-analysis of Randomized Controlled Trials. Sci Rep. 2016;6(1):25325. doi:10.1038/srep25325

21. Nonin. WristOx2 Model 3150 with USB. https://www.nonin.com/support/3150-usb/.

22. Nonin. Nonin Medical’s WristOx2® Model 3150 Beats VirtuOx VPOD in Hypoxia Testing. https://www.nonin.com/resource/wristox2-model-3150-beats-virtuox-vopd-in-hypoxia-testing/.

23. Chen PS, Chiu WT, Hsu PL, et al. Pathophysiological implications of hypoxia in human diseases. J Biomed Sci. 2020;27(1):63. doi:10.1186/s12929-020-00658-7

24. Leach RM, Treacher DF. ABC of oxygen: Oxygen transport---2. Tissue hypoxia. BMJ. 1998;317(7169):1370–1373. doi:10.1136/bmj.317.7169.1370

25. Bhutta BS AFBI. Hypoxia. In: StatPearls [Internet]. Treasure Island (FL): StatPearls Publishing; 2025 Jan-. Available from: https://www.ncbi.nlm.nih.gov/books/NBK482316/.

26. Majumdar SR, Eurich DT, Gamble JM, Senthilselvan A, Marrie TJ. Oxygen Saturations Less than 92% Are Associated with Major Adverse Events in Outpatients with Pneumonia: A Population-Based Cohort Study. Clinical Infectious Diseases. 2011;52(3):325–331. doi:10.1093/cid/ciq076

27. Hurst JH. William Kaelin, Peter Ratcliffe, and Gregg Semenza receive the 2016 Albert Lasker Basic Medical Research Award. J Clin Invest. 2016;126(10):3628–3638. doi:10.1172/JCI90055

28. Elkhenany H, AlOkda A, El-Badawy A, El-Badri N. Tissue regeneration: Impact of sleep on stem cell regenerative capacity. Life Sci. 2018;214:51–61. doi:10.1016/j.lfs.2018.10.057

29. Muscari C, Giordano E, Bonafè F, Govoni M, Pasini A, Guarnieri C. Priming adult stem cells by hypoxic pretreatments for applications in regenerative medicine. J Biomed Sci. 2013;20(1):63. doi:10.1186/1423-0127-20-63

30. Cabarcas SM, Mathews LA, Farrar WL. The cancer stem cell niche—there goes the neighborhood? Int J Cancer. 2011;129(10):2315–2327. doi:10.1002/ijc.26312

31. Pialoux V, Mounier R. Hypoxia-Induced Oxidative Stress in Health Disorders. Oxid Med Cell Longev. 2012;2012:1–2. doi:10.1155/2012/940121

32. Owens RL, Derom E, Ambrosino N. Supplemental oxygen and noninvasive ventilation. European Respiratory Review. 2023;32(167):220159. doi:10.1183/16000617.0159-2022

33. Chourpiliadis C, Bhardwaj A. Physiology, Respiratory Rate.; 2025.

34. Mendo B, Gonçalves M, Lopes L, Matos LC, Machado J. Can Yoga Qigong, and Tai Chi Breathing Work Support the Psycho-Immune Homeostasis during and after the COVID-19 Pandemic? A Narrative Review. Healthcare. 2022;10(10):1934. doi:10.3390/healthcare10101934

35. Russo MA, Santarelli DM, O’Rourke D. The physiological effects of slow breathing in the healthy human. Breathe. 2017;13(4):298–309. doi:10.1183/20734735.009817

36. NIH. Your Guide to Healthy Sleep.; 2011.

37. Zoccoli G, Cianci T, Lenzi P, Franzini C. Shivering during sleep: Relationship between muscle blood flow and fiber type composition. Experientia. 1992;48(3):228–230. doi:10.1007/BF01930460

38. Korthuis R. Exercise Hyperemia and Regulation of Tissue Oxygenation During Muscular Activity. In: Skeletal Muscle Circulation. Morgan & Claypool Life Sciences; 2011. doi:10.4199/C00035ED1V01Y201106ISP023

39. Kump DS. Mechanisms Underlying the Rarity of Skeletal Muscle Cancers. Int J Mol Sci. 2024;25(12):6480. doi:10.3390/ijms25126480

40. Mahler DA, Gifford AH, Waterman LA, Ward J, Machala S, Baird JC. Mechanism of Greater Oxygen Desaturation During Walking Compared With Cycling in Patients With COPD. Chest. 2011;140(2):351–358. doi:10.1378/chest.10-2415

41. Dasanayaka NN, Sirisena ND, Samaranayake N. The effects of meditation on length of telomeres in healthy individuals: a systematic review. Syst Rev. 2021;10(1):151. doi:10.1186/s13643-021-01699-1

42. Carlson BW, Neelon VJ, Carlson JR, Hartman M, Dogra S. Exploratory Analysis of Cerebral Oxygen Reserves During Sleep Onset in Older and Younger Adults. J Am Geriatr Soc. 2008;56(5):914–919. doi:10.1111/j.1532-5415.2008.01672.x

43. Schroeder EC, Franke WD, Sharp RL, Lee D chul. Comparative effectiveness of aerobic, resistance, and combined training on cardiovascular disease risk factors: A randomized controlled trial. PLoS One. 2019;14(1):e0210292. doi:10.1371/journal.pone.0210292

44. Cooper K. Aerobics. 1st ed. Bantam Books; 1968.

45. Cantwell JD. Reflections on Kenneth H. Cooper, MD, MPH. Baylor University Medical Center Proceedings. 2021;34(5):638–639. doi:10.1080/08998280.2021.1930826

